# Parameter Dependence in Identifiability Applied to FP-Fisher-KPP Reaction-Diffusion Equations Parameterized for Tauopathy Network Modeling

**DOI:** 10.1101/2024.10.15.618412

**Authors:** Arsalan Rahimabadi, Habib Benali

## Abstract

Parameterization and a priori identifiability analysis are two interconnected steps that should be carried out in advance of model calibration. In the first place, we propose a framework for parameterizing a recently introduced and analytically studied generalization of the celebrated Fisher-Kolmogorov-Petrovsky-Piskunov (Fisher-KPP) reaction-diffusion (Re-Di) equation with fractional polynomial (FP) terms to model heterogeneous nonlinear diffusion in the propagation of a given species through directed networks, exemplified by the tauopathy progression in Alzheimer’s disease (AD). Next, we present our results on identifiability in a generic sense for the parameterized FP-Fisher-KPP Re-Di equations with regular multi-experimental designs, seamlessly applicable to meromorphic systems, encompassing analytic systems. In particular, the concept of generic local minimal dependence of unknown parameters and regularly parameterized initial conditions will be formalized through the use of one-parameter Lie groups of transformations, and a decomposition method to explore this new concept will be devised. Finally, the Allen Mouse Brain Connectivity Atlas (AMBCA) dataset is utilized to develop a model for tauopathy progression in the mouse brain, which will subsequently be employed to implement the proposed methodology for analyzing a priori identifiability.

## I. Introduction

APRIORI identifiability, in general terms, refers to an inherent characteristic of a dynamical system model that ensures its parameters can be uniquely determined from theoretically available observations, and it has been studied under various definitions and their respective approaches [1], [2]. Differential algebraic approaches are basically available for polynomial and rational dynamical systems to study global and local identifiability by computing input-output relations [3], [4], [5]. However, it has been shown that under certain conditions, Padé and power series approximations may extend results from a differential algebraic setting to test identifiability for systems with non-rational functions [6]. To tackle a priori identifiability in broader classes of systems, including analytic systems, there are typically two main approaches grounded in power series expansions [7], [8], [9] and differential geometry [10], [11], [12]. A key result derived from both approaches is the rank test of the Jacobian matrix of the output vector and its higher-order time derivatives with respect to the unknown parameters and, possibly, initial conditions of a given model.

Without delving into details, local identifiability restricts the uniqueness problem to a neighborhood of a specified vector of unknown parameters in parameter space. Although the rank test serves only as a sufficient condition for examining local identifiability, some previous studies have inaccurately asserted that it is also a necessary condition [8], [9]. As an illustration, it is straightforward to verify that the system 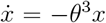 with the output *y* = *x* and a known initial condition *x*_0_ ≠ 0 is locally identifiable at *θ* = 0, while the rank of 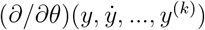 is equal to zero at *θ* = 0 for any nonnegative integer *k*, where *y*^(*k*)^ represents the *k*-th time derivative of *y*. This example can also demonstrate that the reasoning provided at the start of the proof of Theorem 2 in [13] is not generally true, where the conclusion ‘rank(*∂*Φ*/∂θ*) = *q*’ was drawn from the statement ‘the system is *x*_0_-identifiable at *θ* through the input *u*’, and in which *q* denotes the dimension of the parameter vector space associated with *θ* and the mapping Φ incorporates the output *y* and its time derivatives up to the *k*-th degree. We will return to this theorem later.

The concept of local identifiability has generically extended under different terms, such as “structural local identifiability” [14] and “geometric identifiability” [13]. Unlike the definition of structural local identifiability, which does not explicitly incorporate information on initial conditions, geometric identifiability mandates that all initial conditions must be known. Indeed, Xia and Moog [13] adopted distinct definitions, including “identifiability with partially known initial conditions”, to characterize identifiability in a generic sense based on available information regarding initial conditions. Let us highlight the significance of a priori knowledge on initial conditions using the system 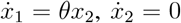 with the output *y* = *x*_1_. It is intuitively evident that the identifiability of the parameter *θ* requires information about *x*_2_. In fact, fixing *x*_2_ to any nonzero real number renders the parameter *θ* locally identifiable at any value of *θ*, implying its identifiability in a generic sense; for further elaboration, see [15], [16], [17]. This example also indicates that the observability-identifiability condition (OIC) [11], [18], where unknown parameters are treated as additional states with zero dynamics and the identifiability problem is reformulated as an observability task, should be approached with caution when the rank deficiency of the observability-identifiability matrix can be compensated by a priori information on initial conditions. Regarding local weak observability, it is important to note that satisfying the observability rank condition (ORC) generically is a necessary and sufficient condition for concluding that a system with smooth functions is locally weakly observable at every point of an open dense subset of the state space, defined as a smooth, connected manifold [19]. In the case of having a priori knowledge about initial conditions, Tunali and Tarn [10] utilized the differential geometry framework of observability introduced in [19] to establish necessary and sufficient conditions for different definitions of local identifiability in certain classes of nonlinear smooth systems. Additionally, Theorems 2 and 4 in [13] present necessary and sufficient conditions for geometric identifiability and for identifiability with partially known initial conditions, respectively, in meromorphic systems based on performing a rank test. However, as discussed in the preceding lines, the reasoning outlined at the beginning of the proof of Theorem 2 is not generally valid, and the proof of Theorem 4 was also left for the readers, perhaps due to its similarity to that of Theorem 2.

Should a test confirm the unidentifiability of a model’s parameters, determining its causes and finding an identifiable reparameterization that preserves the original model’s dynamics and input-output mapping will be the primary focus. Moreover, mechanistic modeling of biological systems frequently results in over-parameterization, which, in turn, gives rise to symmetries in the differential equations of derived models that render their outputs invariant under certain transformations of their parameters, such as scaling [20] and affine types [21]. These symmetries, referred to as Lie symmetries and contributing to unidentifiability, can be investigated using Lie groups of transformations. In general, they can be determined by solving systems of differential equations whose solutions represent the transformations admitted by the models [22], [23], [24]. The reparameterization of a model can also be pursued independently of Lie group theory within the context of differential algebra for polynomial and rational systems [4], [25], or within the framework of power series for systems described by analytic functions [8], [26].

In this manuscript, following Section II, which is devoted to general notations and definitions, Section III delineates a parameterization framework for the extended FP-Fisher-KPP Re-Di equations [27], rooted in a phenomenological perspective of pathology progression in tauopathies, the most prevalent class of neurodegenerative diseases. In Subsection IV-A, we will demonstrate the sufficiency and necessity of a generic rank test as well as the existence of identifiable reparameterizations for the proposed parameterized FP-Fisher-KPP Re-Di equations with regular multi-experimental designs in a power series setting, which are readily extendable to meromorphic systems with regularly parameterized initial conditions. Reflecting on our literature survey, it is important to emphasize that we found no concrete proof in prior manuscripts supporting the necessity of a rank test defined for analytic or meromorphic systems with fully or partially known initial conditions in a generic sense. Furthermore, our result on identifiable reparameterizations is a generic refinement of Theorem 1 in [8]. Afterward, Subsection IV-B is dedicated to introduce the concept of generic local minimal dependence of unknown parameters and regularly parameterized initial conditions using one-parameter Lie groups of transformations and to propose a decomposition method to analyze this new concept. Eventually, in Section V, a tauopathy progression model in the mouse brain is constructed using the Allen Mouse Brain Connectivity Atlas (AMBCA) dataset, which will then be employed to apply the proposed identifiability framework.

## II. General notations and definitions

The set of nonnegative integers and the set of the first *N* positive integers are represented by 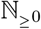 and ℕ_≤*N*_, respectively. Here, ℝ^*M*^, 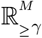, and 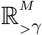 denote the sets of all *M* -tuples whose components belong to (−∞, ∞), [*γ*, ∞), and (*γ*, ∞), respectively, given *γ* ∈ (−∞, ∞). The set of all *M*_1_-by-*M*_2_ matrices whose entries belong to [0, ∞) is indicated by 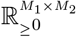, and the maximum absolute column sum of the matrix is represented by || · ||_1_. Furthermore, let 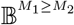 be the set of full-rank *M*_1_-by-*M*_2_ binary matrices, with *M*_1_ ≥ *M*_2_, having exactly one entry of 1 in each row and zero elsewhere. Throughout the paper, boldface is used to distinguish vectors and matrices from scalars. Suppose ***u, v, w*** ∈ ℝ^*M*^, and let **Λ** (***v***) be a diagonal matrix whose diagonal elements are the elements of ***v***. Then, the vector ***v*** ⊙ ***w*** defined by ***v*** ⊙ ***w*** = **Λ** (***v***) ***w*** is called the Hadamard product of ***v*** and ***w*** [28]. The set of all elements of ***v*** is also symbolized by {{***v*** }}, and inversely, vec ({{***v***}}) denotes the vectorization of all elements of {{***v***}}. Moreover, we define functions 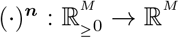 by 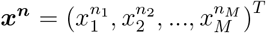 where 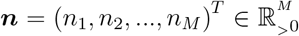 and

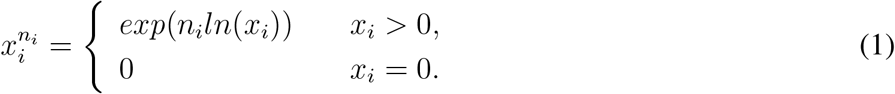

It is obvious that ***x***^***n***^ = **Λ** (***x***^***n***^) **1**_*M*_ where **1**_*M*_ stands for the *M* -dimensional all-ones vector. Lastly, consider the equivalence relation (*f, g*) ∼ (*f* ^′^, *g*^′^) whenever *fg*^′^ = *gf* ^′^ on the set of all pairs (*f, g*) of elements of the ring of analytic functions from a connected open subset of a Euclidean space of a given dimension to ℝ, such that *g* ≠ 0. The equivalence class of (*f, g*) is denoted by *f/g*, and the elements of the corresponding quotient field are called meromorphic functions. Accordingly, a vector-valued function is said to be meromorphic if each of its component functions is a meromorphic function.

## III. Model description and parameterization

Here, we focus on parameterizing FP-Fisher-KPP Re-Di equations for modeling species propagation within neuronal networks.

### A. Phenomenological insights into tauopathy progression

Discovered in 1975, tau protein contributes substantially to the stability and flexibility of microtubules, which form the backbone of axons and dendrites and maintain brain connectivity. Despite its unfolded form enhancing hydrophilicity, hyperphosphorylation and truncation can promote tau aggregation [29], characterizing tauopathies, a major category of over 26 identified neurodegenerative diseases, among which Alzheimer’s disease (AD) solely accounts for 60–80% of dementia cases [30]. Injecting tau seeds into the mouse brain induces tau pathology, supporting a prion-like hypothesis under which misfolded tau species recruit normal tau monomers to form larger aggregates, facilitating the spread of pathology from the injection sites to synaptically connected brain regions [31], [32]. Indeed, the propagation of toxic tau species is predominantly attributed to axonal transportation [33]. Although experimental evidence confirms the trans-synaptic transmission of misfolded tau species between neurons in both anterograde and retrograde directions [34], [35], a recent study suggests that tau pathology propagates more efficiently in the retrograde direction [36], [37], an important observation has been overlooked in works utilizing undirected graph models for neuronal networks in investigating tauopathy progression [38], [39], [40], [41]. In addition, given that axonal bundles have fibrous structures that create porous media, it is expected that tau pathology spreads through a nonlinear diffusion process at the mesoscale. This aspect has not been addressed in recent studies employing linear Laplacian operators to model the spatiotemporal evolution of tauopathies [36], [38], [40], [41], [42], [43]. The heterogeneous morphology of neuronal networks also gives rise to the hypothesis that the properties of nonlinearity and directionality in tauopathy progression may vary across different regions of the brain.

Drawing from the aforementioned description of tauopathies, Fig. 1 illustrates a schematic topological representation of two mutually connected neuronal populations. Species can typically spread through each connection in both anterograde and retrograde directions with different transmission rates. Additionally, each population has its own species-specific characteristics, leading to a distinct capacity to localize the spread of a given species. This general concept of neuronal networks will be employed to parameterize the equations.

**Fig. 1:**
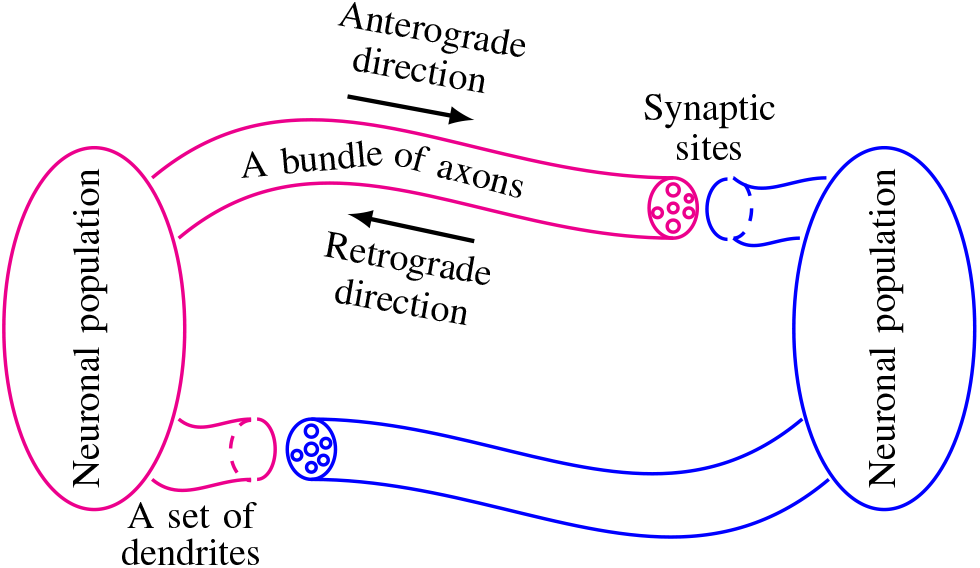
A schematic topological representation of two neuronal populations that are mutually connected.

### B. A generalized directed graph representation for neuronal networks

A node-capacitated bi-weighted directed graph (or simply an NCB digraph) *G* with no self-loops is a quintuple (*N*_*G*_, *E*_*G*_, *ω*_*G*_, 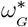, *n*_*G*_), where *N*_*G*_ is a finite nonempty set of nodes (or vertices), *N*_*G*_ = ℕ_≤*M*_ with a positive integer *M*, *E*_*G*_ is a set of directed edges (or links), *E*_*G*_ ⊆ *N*_*G*_ × *N*_*G*_ \ {(*k, i*) ∈ *N*_*G*_ × *N*_*G*_ : *k* = *i*}, *ω*_*G*_ : *E*_*G*_ → [0, ∞) and 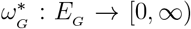 are functions that associate each edge of *E*_*G*_ from vertex *k* to vertex *i, k* → *i*, to an anterograde strength *ω*_*ik*_ and to a retrograde strength 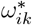, respectively, such that max (*ω*_*ik*_, 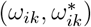) *>* 0 (if there is no such edge, we set 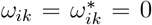), and *n*_*G*_ : *N*_*G*_ → [1, ∞) is a function that assigns a capacity exponent *n*_*i*_ to each node *i*. Given that ***W*** _*G*_ = [*ω*_*ik*_] and 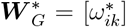 represent the anterograde strength adjacency (ASA) and the retrograde strength adjacency (RSA) matrices of G, respectively, the corresponding digraph of *G* is naturally defined by identifying 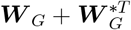 as its weighted adjacency matrix. Accordingly, we say that *G* is strongly connected if its corresponding digraph is strongly connected. For information regarding the strong connectivity properties of digraphs, the interested reader is referred to the subsection 2.3 of [27].

### C. Derivation of a parameterized diffusion model

(*Transmission rate law*) For a given link *k* → *i* of a NCB digraph *G* with *M* nodes, it is here postulated that the transmission rate of a given species can be described by

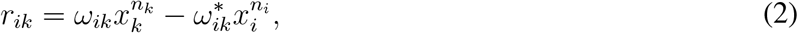

which may be interpreted as a generalization of Fick’s first law of diffusion, relating the diffusive flux to the gradient of the concentration, in a graph framework. (*Conservation of mass*) To uphold the conservation of mass for diffusive species, the transmission rates must conform the following equations:

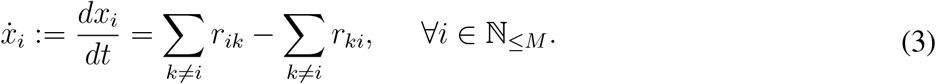

Given ***W*** _*G*_ = [*ω*_*ik*_], 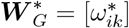, ***n*** = (*n*_1_, *n*_2_, …, *n*_*M*_)^*T*^, and ***x*** = (*x*_1_, *x*_2_, …, *x*_*M*_)^*T*^, Eq.(3) can be rewritten as

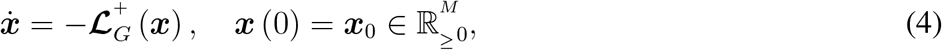

with

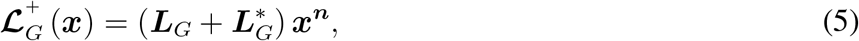

where

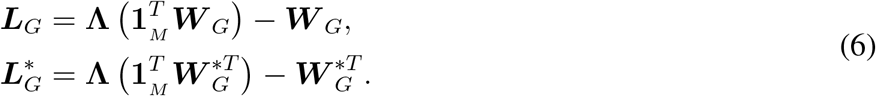

Note that the function 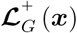technically represents the restriction of the function ℒ_*G*_ (***x***) introduced in [27] to the nonnegative orthant.

In practice, an essential condition for Eq.(4) to accurately describe a diffusion process is to use the matrices ***W***_*G*_ and 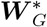 which are well-estimated based on not only the morphological characteristics of the given neuronal network, such as the length and diameter of axonal fibers, but also its species-specific characteristics, including the mobility and cell-to-cell transmissibility of the given species. Thus, in order to be able to separately investigate the effects of its morphological and species-specific characteristics on diffusion, we adopt the following parameterizations of the ASA and RSA matrices:

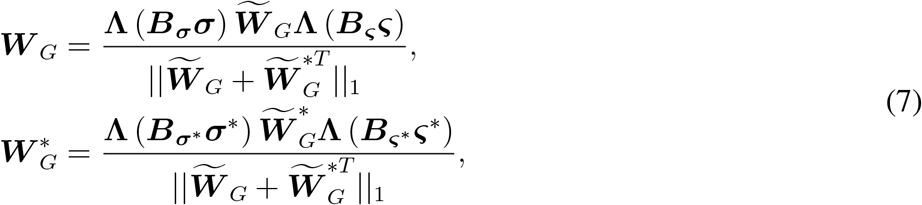

where 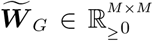 and 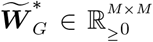 are here termed the morphological ASA and RSA matrices, respectively, that are computed using the morphological characteristics of the given neuronal network. Further, 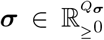 and 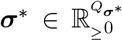, with *Q*_***σ***_, 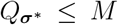, denote the anterograde and the retrograde target-dependent diffusion parameter vectors, respectively, and 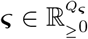 and 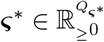, with *Q*_***ς***_, 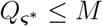, represent the anterograde and the retrograde source-dependent diffusion parameter vectors, respectively. The parameter vectors ***σ*** and ***σ***^*^ are said to be target-dependent for the reason that each row of the morphological ASA and RSA matrices, corresponding to the directed edges with the same target node, is multiplied by an element of these vectors, and similarly, ***ς*** and ***ς***^*^ are said to be source-dependent since each column of the morphological ASA and RSA matrices, which corresponds to the directed links with the same source node, is multiplied by an element of ***ς*** or ***ς***\*. Moreover, 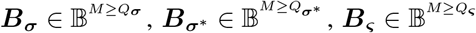, and 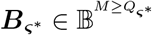 are referred to as nodal parameter distribution (NPD) matrices because they incorporate information on how the elements of the parameter vectors are assigned to the nodes of *G*.

Given a parameter vector 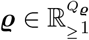 and its NPD matrix 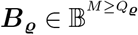, the capacity exponent vector ***n*** in Eq.(5) may also be parameterized as

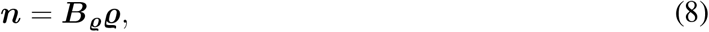

and the initial condition of Eq.(4) can be written as

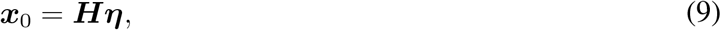

where 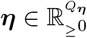 and 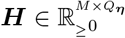 with rank *Q*_***η***_.

### D. A parameterized FP-Fisher-KPP Re-Di model

To model a reaction-diffusion process of a given species on a NCB digraph *G* with *M* nodes, we introduce a modification to Eq. (4) by incorporating a parameterized reaction term as follows:

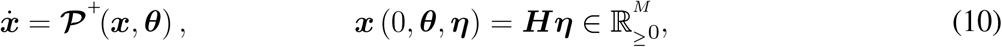

with

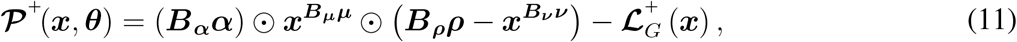

where the reaction coefficient parameter vectors 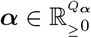 and 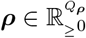 and the reaction exponent parameter vectors 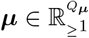 and 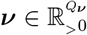 have their own NPD matrices 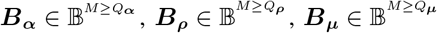 and 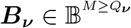, respectively. Given the concatenated parameter vector

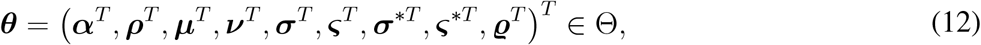

with the Cartesian product

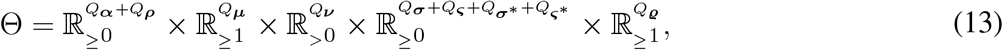

and the concatenated NPD matrix

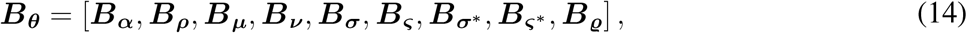

the pair (***θ, B***_***θ***_) is called the parameterization of the model (10). Note that the Cartesian product is here defined such that its elements are represented as column vectors. Regardless of the chosen parameterization, the model (10) technically falls within the class of (extended) FP-Fisher-KPP Re-Di equations, thereby possessing the following property.

#### Property 3.1

*(Positivity and boundedness [27]):* The initial value problem (10) has a unique solution ***x*** (*t*, ***θ, η***) remaining entirely in a compact subset of 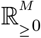 for all *t* ≥ 0. Moreover, strong connectivity of *G* can guarantee the existence of strictly positive solutions.

#### Property 3.2

*(Strict positivity):* Assume that *G* is strongly connected. For any 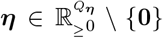, the system (10) has a unique solution ***x*** (*t*, ***θ, η***) that lies entirely in 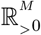 for all *t >* 0.

*Proof:* See Appendix A .

## IV. A priori identifiability and reparameterization

Now, let us take into account a set of *J* experiments on the model (10) with the initial concentration vectors ***x***_*j*_ (0, ***θ, η***_*j*_) = ***H***_*j*_***η***_*j*_, where 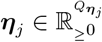 and 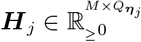 with rank 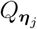, and the measurement vectors ***y***_*j*_ = ***h***_*j*_(***x***_*j*_, ***θ***), where ***h***_*j*_(***x***_*j*_, ***θ***) is a 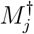-dimensional vector-valued meromorphic function of 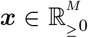 and ***θ*** ∈ Θ. A theoretical design (or simply a design) for these experiments is specified as a collection comprising:

- The parameterization (***θ, B***_***θ***_) ;
- The set of all pairs (***η***_*j*_, ***H***_*j*_); and
- The set of all functions ***h***_*j*_ (***x***_*j*_, ***θ***), *j* ∈ ℕ_≤*J*_ .

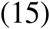

Let us also define a vector ***ϑ*** of minimal dimension encompassing all unknown parameters and unknown initial conditions; thus,

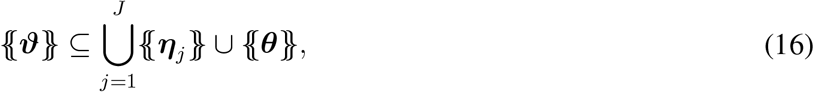

and the Cartesian product of the prescribed ranges of its elements is denoted by Θ_***ϑ***_. Accordingly, the knowns can be specified by

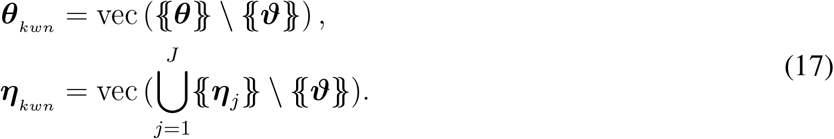

In this section, we aim to investigate the possibility of uniquely determining a vector of unknowns ***ϑ*** confined to Θ_***ϑ***_, considering the design (15).

### A. Multi-experimental generic local identifiability

Given the model (10) with the design (15), a vector of unknowns ***ϑ*** is said to be generically locally (g.l.) identifiable if there exists an open and dense subset 𝔇 of the Cartesian product 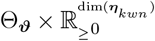 such that for any vector 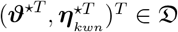, there exists a neighborhood 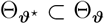 containing ***ϑ***^⋆^ such that for any two ***ϑ***_1_, 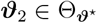 and any *t*^⋆^ *>* 0 satisfying

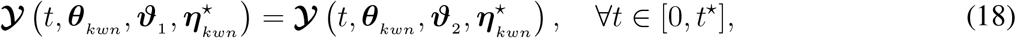

Where

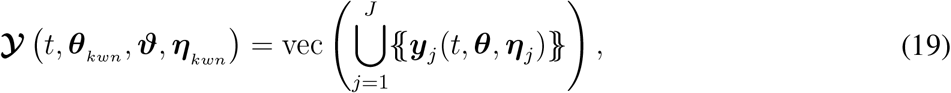

it can be concluded that ***ϑ***_1_ = ***ϑ***_2_ . Otherwise, it is said to be g.l. unidentifiable. This definition extends the conventional structural identifiability [10], [13] to a multi-experimental setup, where 𝒴 (*t*, ***θ***_*kwn*_, ***ϑ, η***_*kwn*_) can be regarded as the measurement vector of the augmented system with the augmented state vector 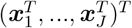, constructed by treating all *J* experiments as a single comprehensive experiment.

Thanks to the analyticity of the right-hand side of Eq.(10) in 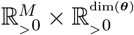, the function ***y***_*j*_ (*t*, ***θ, η***_*j*_) is analytic in 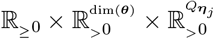, and thus, it can be locally described by a Maclaurin series with respect to *t*, whose *i*-th coefficient is expressed as:

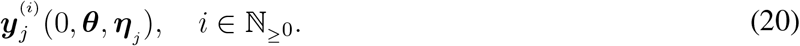

Therefore, we can recast the given condition for the generic local identifiability as follows: ***ϑ*** is g.l. identifiable if and only if for any vector 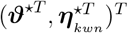 belonging to an open dense subset 𝔇 of 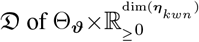, the infinite set of the equations

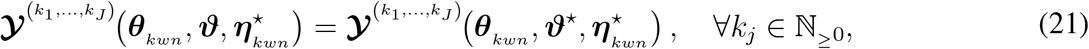

where

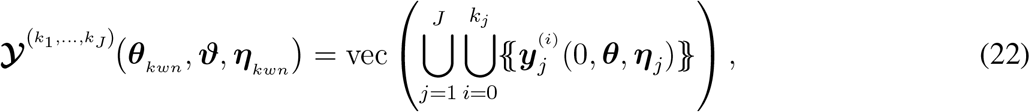

has the unique solution ***ϑ*** = ***ϑ*** in a neighborhood 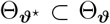 containing ***ϑ***. This new condition is a multi-experimental generic version of the necessary and sufficient condition originally provided in [7].

#### Property 4.1

*(Generic rank):* The rank of 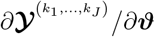 is constant on an open dense subset 𝔇 of 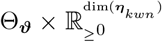, arising from a fundamental result that the zero set of a non-identically zero analytic function defined on a connected open subset of a Euclidean space of a given dimension has an empty interior. In fact, it makes sense to define the generic rank of 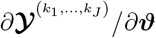 as the dimension of the maximum square submatrix having non-identically zero determinant, which coincides with its rank at the points of 𝔇. It is obvious that its generic rank is greater than or equal to its rank at any point of the interior of 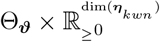.

#### Property 4.2

*(Adequate degree of differentiation [44], [9]):* Consider the augmented system

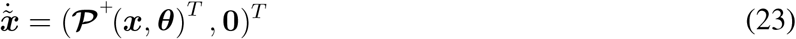

with the augmented state vector 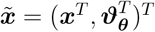, where ***ϑ***_***θ***_ exclusively comprises the unknown parameters, consequently, {{***ϑ***_***θ***_}} ⊆ {{***θ***}}, and a meromorphic function *y* of 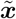. It can be shown that if, for some 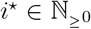, the row vector 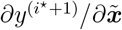 is linearly dependent on the set of 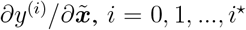, *i* = 0, 1, …, *i*^⋆^, over the field of meromorphic functions of 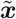, then the row vector 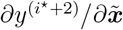 is also linearly dependent on the same set, where *y*^(*i*)^ represents the *i*-th time derivative of *y*. Hence, since a single nonzero row vector is linearly independent of itself and the generic rank of 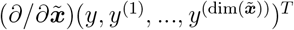 is less than or equal to 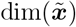, the smallest such nonnegative integer *i*^⋆^ exists and is less than or equal to 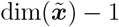. Consistently, given the system (23) with a vector-valued meromorphic function ***y*** of 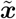, one can establish that if, for some 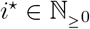, the generic rank of

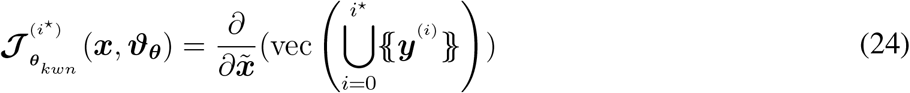

is equal to the generic rank of the augmented Jacobian 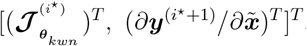, then any augmented Jacobian including the derivatives of degrees higher than *i*^⋆^ has the same generic rank. The smallest such nonnegative integer *i*^⋆^ is here called the adequate differentiation degree with respect to ***ϑ***_***θ***_ (ADD-***ϑ***_***θ***_) of s***y***, which is similarly bounded above by

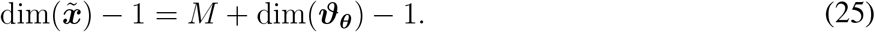

Example 4.1 highlights that Property 4.2 is a generic property, rather than being point-specific; let us reiterate that generic properties are of interest in this manuscript.

#### Example 4.1

*([17]):* Consider the augmented system 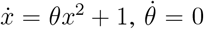 with the output *y* = *x* and the specific initial condition *x*(0) = 0. Although the rank of the Jacobian matrix 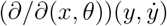 is equal to 1 for all *θ* ∈ (−∞, ∞) and *x* = 0, the rank of 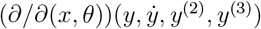 equals 2 when *x* = 0.

In the case of parameterized initial conditions, Property 4.2 holds provided that the following property is satisfied.

#### Property 4.3

*(Regularly parameterized initial conditions):* The initial condition ***x*** (0, ***θ, η***) = ***Hη*** is said to be regularly parameterized if the generic ranks of 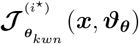 in (24) and 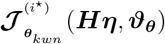 are identical, where *i*^⋆^ is equal to ADD-***ϑ***_***θ***_ of ***y***. Accordingly, the design (15) is called regular if all the initial conditions are regularly parameterized.

#### Example 4.2

Consider the system 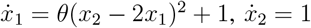 with the parameter *θ* and the output *y* = *x*_1_. For the parameterized initial condition (*x*_1_(0, *θ, η*), *x*_2_(0, *θ, η*)) = (*η*, 2*η*), it can be shown that Property 4.3 fails to hold, and one can check that while the generic rank of 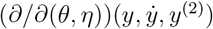 equals 1, the generic rank of 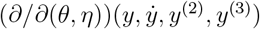 is equal to 2. On the other hand, it can be verified that the initial condition (*η, η*) is regularly parameterized.

The implicit function theorem, along with the (constant) rank theorem, offers an analytical approach to investigating the uniqueness of the solution to the infinite set of Eq.(21), as delineated in the following test.

#### Test 4.1

*(Multi-experimental generic local identifiability):* Given the model (10) with the regular design (15) (resp., the design (15)), a vector of unknowns ***ϑ*** is g.l. identifiable if and only if (resp., if)

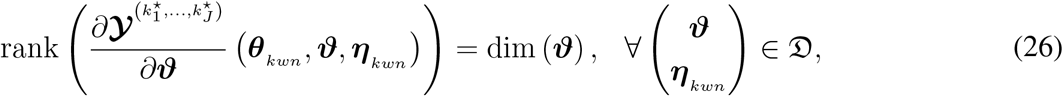

where 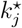 stands for the ADD-***ϑ***_***θ***_ of ***y***_*j*_ (resp., 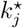 can be any nonnegative integer), and 𝔇 denotes an open dense subset of 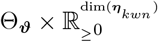.

*Proof:* (Sufficiency) Given a vector 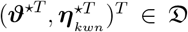, and knowing that there is at least one solution ***ϑ*** = ***ϑ***^⋆^ to the infinite set of Eq. (21), we only need to locally establish its uniqueness. It follows promptly from the condition (26) that there is at least one full-rank dim (***ϑ***)-by-dim (***ϑ***) submatrix of 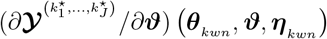 whose corresponding subvector of 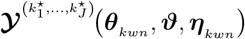 meets the conditions of the implicit function theorem [45], which locally guarantees the uniqueness of the solution ***ϑ*** = ***ϑ***^⋆^ to the infinite set of Eq. (21).

(Necessity) To prove the necessity, we aim to use a contradiction argument. Suppose that the generic rank of 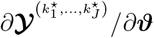 is equal to λ ∈ {0} ∪ ℕ_≤dim(***ϑ***)−1_. Note that for *λ* = 0, the proof is trivial. For a vector 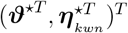 at which the rank of 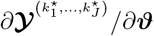 equals *λ* ∈ ℕ_≤dim(***ϑ***)−1_, it follows from the rank theorem [45] that there exist neighborhoods 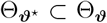 of ***ϑ***^⋆^ and 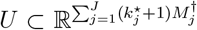 containing 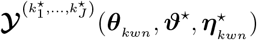 such that

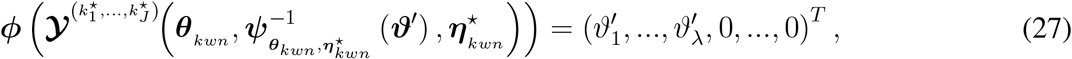

where 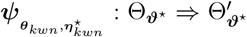 and ***ϕ*** : *U* ⇒ *U*^′^ are diffeomorphisms with open subsets 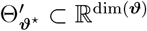 and 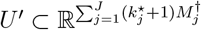 Supposing that ***ϕ***^⋆^ represents the last 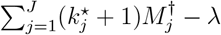 components of ***ϕ***, we have

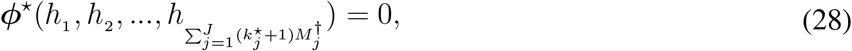

where *h*_*l*_ represents the *l*-th element of 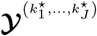. Thanks to the invertibility of ***ϕ***, the Jacobian matrix 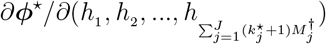 must have a rank equal to 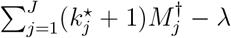. Therefore, it follows from the implicit function theorem that Eq. (28) can be locally solved for 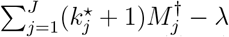 number of *h*_*l*_ ‘s as functions of the remaining *λ h* ‘s, denoted by 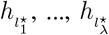. Hence, the function 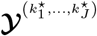 can be locally reparameterized by

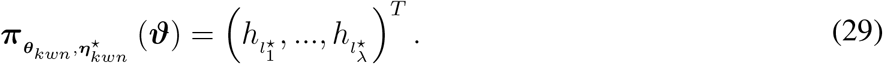

Now, consider a nonnegative integer vector (*k*_1_, …, *k*) for which the rank of 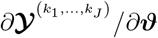 is equal to *λ* at 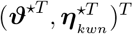. Similar to our approach for obtaining 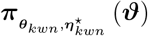 from 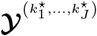, it can be shown that there is also a local reparameterization

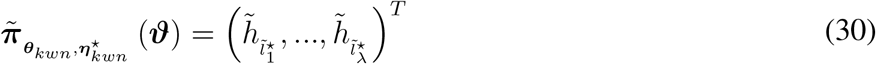

for 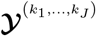, where 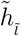 indicates its 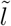-th element. Consequently, it can be inferred from Lemma 4.1 that the function 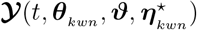 can be locally reparameterized by 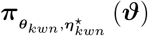; that is to say that there locally exists a function 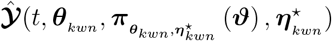 such that

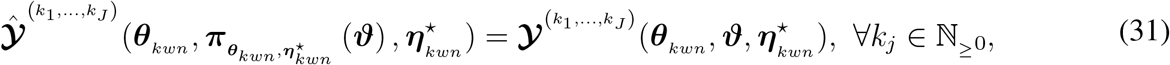

which locally implies

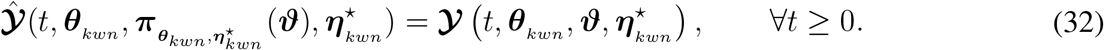

It can be deduced from the preimage theorem, also known as the regular level set theorem [45], that the preimage of 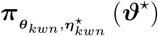 under 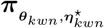 is a smooth manifold of dimension (dim(*ϑ*) − *λ*) *>* 0; thus, in addition to ***ϑ***^⋆^, there are infinitely many vectors ***ϑ*** in the preimage of 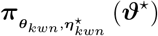 that are contained in any neighborhood of ***ϑ***^⋆^, which contradicts the local uniqueness of ***ϑ***^⋆^.

#### Lemma 4.1

Consider the reparameterizations (29) and (30). There locally exist analytic functions ***T*** and 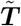 such that

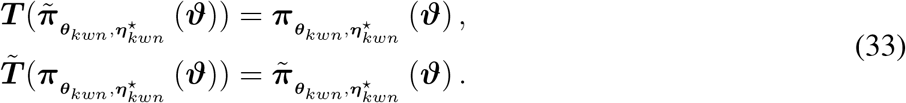

*Proof:* See Appendix B .

In light of the argument provided for the necessity of Test 4.1, particularly Lemma 4.1, we can draw the following conclusion.

#### Proposition 4.1

*(G.l. identifiable reparameterizations):* Consider the model (10) with the regular design (15), and assume that the Jacobian 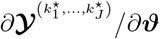 has generic rank *λ* ∈ ℕ_≤dim(***ϑ***)_. The function 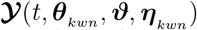 can be generically locally reparameterized in terms of any subset comprising *λ* elements of 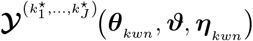, denoted by 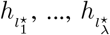, if the generic rank of 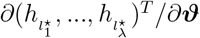 is equal to λ.

### B. Generic local minimal dependence decomposition of the set of unknowns

In this part, the origins of the generic local unidentifiability are explored by defining the concept of generic local minimal dependence of unknowns using (one-parameter) Lie groups of transformations. Subsequently, we will introduce a method for obtaining subsets of unknowns that are generically locally (g.l.) minimally dependent and for determining a subset with the minimal number of unknowns such that the remaining ones are g.l. identifiable.

Consider the model (10) with the regular design (15). Let us first rewrite the functions (19) and (22) as follows:

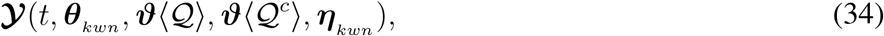

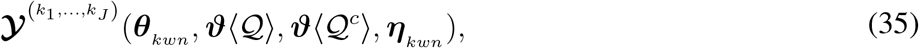

respectively, where ***ϑ***⟨𝒬⟩ and ***ϑ***⟨𝒬^*c*^⟩ denote subvectors of the vector of unknowns ***ϑ*** consisting of elements whose indices belong to a nonempty set 𝒬 ⊆ ℕ_≤dim(***ϑ***)_ and its complement 𝒬^*c*^ = ℕ_≤dim(***ϑ***)_ \ 𝒬, respectively, sorted in ascending order according to their indices. Further, let Θ_***ϑ***⟨𝒬⟩_ represent the Cartesian product of the prescribed ranges of the elements of ***ϑ***⟨𝒬⟩. Given an open set *O* ⊂ Θ_***ϑ***⟨𝒬⟩_ and an interval *I* ⊆ ℝ including zero, the set of transformations

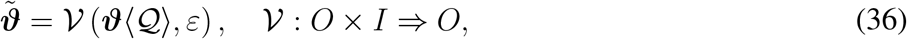

depending on the parameter *ε*, forms a (one-parameter) Lie group of transformations on *O* [46] if

- For each *ε* ∈ *I*, the transformations are one–to–one onto *O*;
- 𝒱 is smooth with respect to *ϑ*⟨𝒬⟩ in O;
- 𝒱 is an analytic function of *ε* in I;
- 𝒱 (*ϑ*⟨𝒬⟩, 0) = *ϑ*⟨𝒬⟩; and
- 𝒱 (𝒱 (*ϑ*⟨𝒬⟩, *ε*_1_), *ε*_2_) = 𝒱 (*ϑ*⟨𝒬⟩, *ε*_1_ + *ε*_2_), ∀*ϑ*⟨𝒬⟩ ∈ *O*, ∀*ε*_1_, *ε*_2_ ∈ *I*.

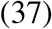

For sufficiently small |*ε*|, 𝒱 (***ϑ***⟨𝒬⟩, *ε*) can be expanded in a Maclaurin series to obtain

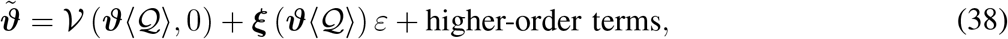

Where

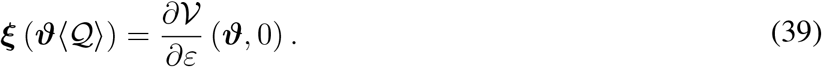

The elements of ***ξ*** (***ϑ***⟨𝒬⟩) are referred to as the infinitesimals of the Lie group of transformations (36).

#### Theorem 4.1

*(Lie’s first fundamental theorem [46]):* The Lie group of transformations (36) is equivalent to the solution of the initial value problem for the system of differential equations

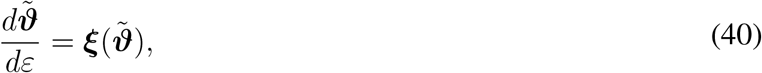

with

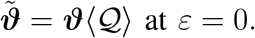

For a vector 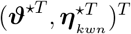 belonging to the interior of 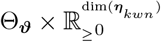, the function

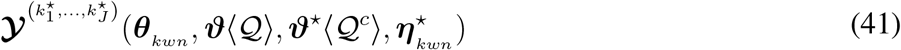

is said to be invariant under the Lie group of transformations (36) if and only if

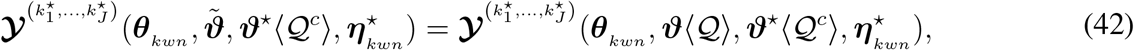

for any ***ϑ***⟨𝒬⟩ ∈ *O* and *ε* ∈ *I*.

#### Theorem 4.2

*(Invariant functions [46]):* The function (41) is invariant under the Lie group of transformations (36) if and only if

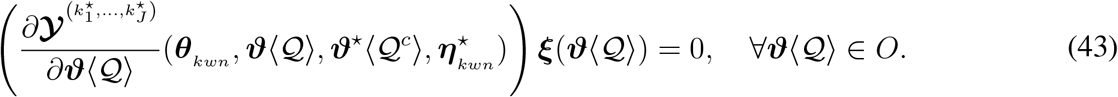

#### Corollary 4.1

If the function (41) is an invariant function under the Lie group of transformations (36), then

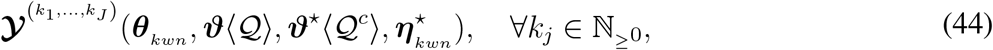

is also invariant under (36), ensuring that the function (34), with ***ϑ***^⋆^⟨𝒬^*c*^⟩ and 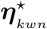, remains locally invariant under (36).

*Proof:* It is immediately deduced from Theorem 4.2 and the fact that each row of 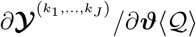 can be written as a linear combination of a set of rows of 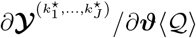.

Now, we are going to define the generic local minimal dependence of unknowns. The elements of ***ϑ***⟨𝒬⟩ are said to be generically locally (g.l.) minimally dependent if there exists an open dense subset 𝔇 of 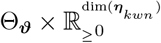 such that, for any vector 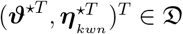, there exists a Lie group of transformations

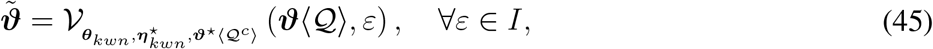

defined in a neighborhood 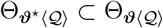 containing ***ϑ*** ⟨𝒬⟩, satisfying the following conditions:

- All of its infinitesimals are non-vanishing in 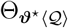;
- The function (41) remains invariant under it; and
- The set 𝒬 is minimal with respect to the function (41); i.e., no nonempty proper subset 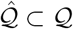 exists such that the function 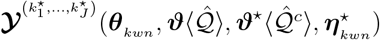 admits a nontrivial Lie group of transformations.

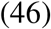

Let 𝕍^*𝔡*^ denote the vector space of all *𝔡*-dimensional vector-valued meromorphic functions of ***ϑ*** and ***η***_*kwn*_, which are also dependent on ***θ***_*kwn*_, over the field of meromorphic functions of ***ϑ*** and ***η*** _*kwn*_, which are also dependent on ***θ***_*kwn*_. A set of vectors ***u***_1_, …, ***u***_*i*⋆_ from a vector space 𝕍^*𝔡*^ is said to be minimally linearly dependent if there exist non-identically zero meromorphic functions *γ*_*i*_ such that 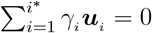, and no nonempty proper subset of the vectors is linearly dependent. Next, take into account the function (35) with 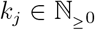 for all *j*. As a complement to the generic rank of 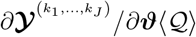, we can also define its generic nullity which is equal to the subtraction of its generic rank from dim(***ϑ***⟨𝒬⟩). Moreover, its kernel on 𝕍^dim(***ϑ***⟨𝒬⟩)^ can be defined as follows:

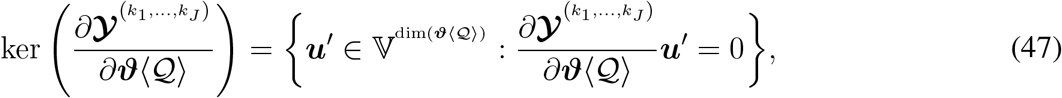

whose dimension equals the generic nullity of 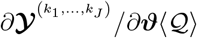. It is evident that the generic nullity of 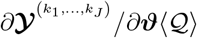 is lower than or equal to its nullity at any point of the interior of 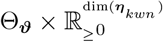. Additionally, the following lemma can be inferred straightforwardly.

#### Lemma 4.2

Given the function (35) with 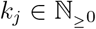 for all *j*, the columns of 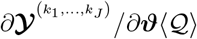 are minimally linearly dependent if and only if its generic nullity is equal to one and its kernel given by (47) contains a vector-valued meromorphic function whose elements are all non-identically zero.

#### Proposition 4.2

*(Generic local minimal dependence):* Considering the function (35) with *k*_*j*_ = *k*^⋆^ for all *j*, the elements of ***ϑ***⟨𝒬⟩ are g.l. minimally dependent if and only if the columns of 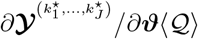 are minimally linearly dependent.

*Proof:* (Sufficiency) Owing to the fact that the columns of 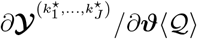 are minimally linearly dependent, Lemma 4.2 directly implies that its generic nullity is equal to one and its kernel given by (47) has a vector-valued meromorphic function ***ξ***(***θ***_*kwn*_, ***ϑ***⟨𝒬⟩, ***ϑ***⟨𝒬^*c*^⟩, ***η***_*kwn*_) whose elements are all non-identically zero. Hence, there is an open dense subset 𝔇 of 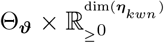 such that, for any vector 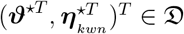, we have

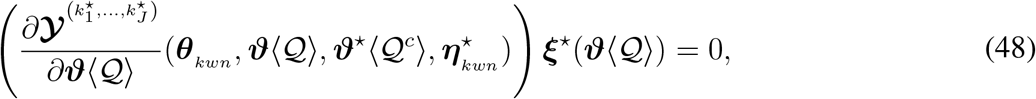

with

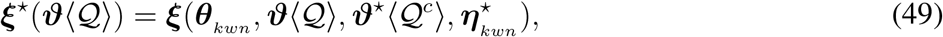

defined in a neighborhood 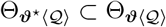 containing ***ϑ*** ⟨𝒬⟩, such that

- All of its elements are non-vanishing in a neighborhood 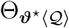; and
- nullity 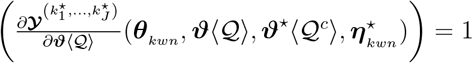 for all 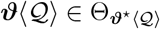.

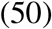

Consequently, it follows from Theorems 4.1 and 4.2 that there exists a Lie group of transformations whose vector of infinitesimals is ***ξ***^⋆^(***ϑ***⟨𝒬⟩), satisfying the first two conditions given in (46). The last condition, namely that 𝒬 is minimal, can be shown by contradiction. Suppose that there is a nonempty proper subset 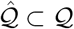 such that the function 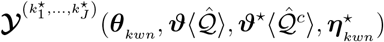 admits a nontrivial Lie group of transformations whose vector of infinitesimals is denoted by 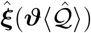, defined in a neighborhood 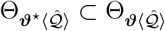 containing 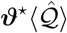. Thus, Theorem 4.2 implies

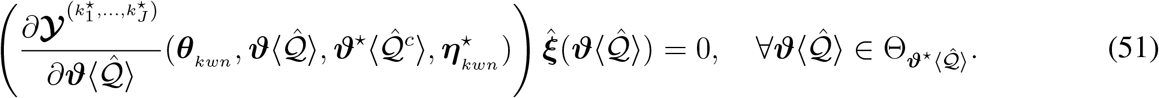

On the other hand, since 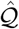 is a nonempty proper subset of 𝒬, it is deduced from the properties given in (50) that there must be a neighborhood 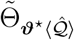 containing 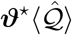 such that

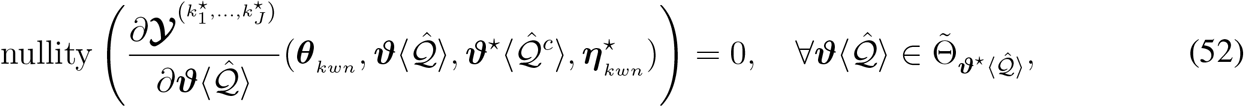

which contradicts the equality (51).

(Necessity) By definition, it follows from the conditions given in (46) and Theorem 4.2 that there exists an open dense subset 𝔇 of 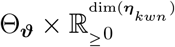 such that, for any vector 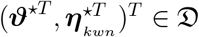,

- There is a vector 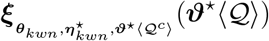 with all nonzero elements; and
- nullity 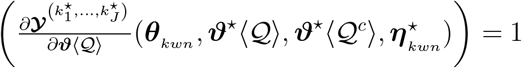;

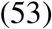

where 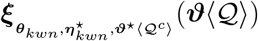 indicates the vector of the infinitesimals of the Lie group of transformations corresponding to 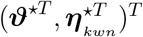. Moreover, Let 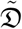 denote the open dense subset of 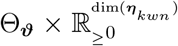 on which the rank of 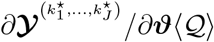 remains constant. Due to the fact that the intersection of two open dense subsets of 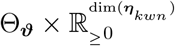 is also an open dense subset thereof (as per the Baire category theorem [47]), it is deduced from the conditions given in (53) that the generic nullity of 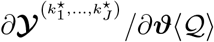 is equal to one, and there is also a function defined by

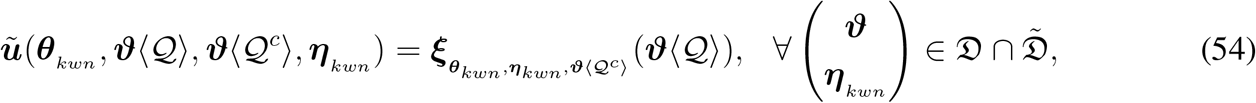

such that all of its elements are non-vanishing in 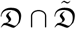, and

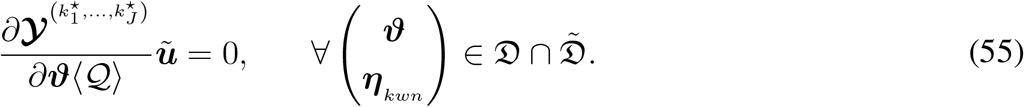

Note that, in general, we cannot yet assert that *ũ* belongs to 𝕍^dim(***ϑ***⟨𝒬⟩)^ . Owing to the fact that the generic nullity of 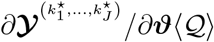 is equal to one, its kernel on 𝕍^dim(***ϑ***⟨𝒬⟩)^ can be spanned by only a vector-valued meromorphic function *û*. For some ℝ-valued function 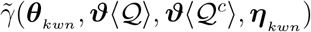, we must have

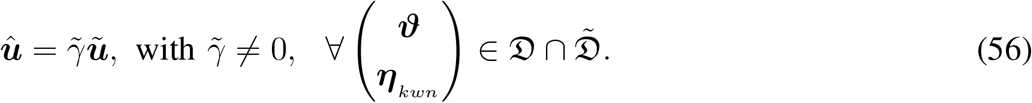

As a result, since all elements of *ũ* are non-vanishing in 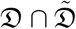, it is inferred that the elements of *û* are non-identically zero, and by Lemma 4.2, we can conclude that the columns of 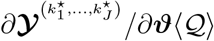 minimally linearly dependent.

◼

Before stating the next result, we would like to specify the reduced column echelon form (RCEF) of a matrix over the field of meromorphic functions of given arguments, analogous to that of a matrix over ℝ [28]. The defining characteristics of the RCEF are as follows:

1. All identically zero columns are on the right;
2. The leading entry of any non-identically zero column is a 1;
3. All other entries in the row of a leading entry are zero; and
4. If *l*_1_ *> l*_2_, then the leading one in the *l*_1_-th column appears below that in the *l*_2_-th column;

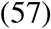

where the leading entry of a non-identically zero column is its first non-identically zero element when read from top to bottom. Furthermore, the RCEF of a matrix can be uniquely obtained by performing a finite sequence of elementary column operations: (I) interchange of two columns, (II) multiplication of a column by a non-identically zero meromorphic function, and (III) addition of a scalar multiple of one column to another column.

#### Proposition 4.3

*(Minimal dependence decomposition):* Consider the model (10) with the regular design v(15), and assume that the rank of the Jacobian 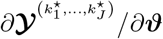 is equal to *λ* ∈ {0} ∪ ℕ_≤dim(***ϑ***)−1_ on an open dense subset 𝔇 of 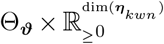, and let *λ*^*c*^ denote its generic nullity; thus, *λ*^*c*^ = dim(***ϑ***) − *λ*. Moreover, designate 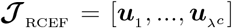 as the RCEF of a dim (***ϑ***)-by-*λ*^*c*^ matrix whose columns spanning the kernel of 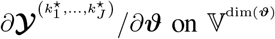 on 𝕍^dim(***ϑ***)^. Let 𝒬_*l*_ and 𝒬 _*LE*_ also represent the sets of indices corresponding to non-identically zero elements of ***u***_*l*_ and to the rows of ***𝒥***_RCEF_ including the leading entries of its columns, respectively. Then, the following hold:

1. For every 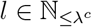, the elements of ***ϑ***⟨𝒬_*l*_ ⟩ are g.l. minimally dependent;
2. For any fixed subvector 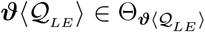 such that 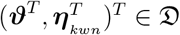 (i.e., for almost all chosen values of ***ϑ***⟨𝒬 _*LE*_ ⟩), the subvector of the remaining unknown ***ϑ***⟨ℕ_≤dim(***ϑ***)_ \ 𝒬 _*LE*_ ⟩ becomes g.l. identifiable; and
3. The set 𝒬 _*LE*_ includes the minimal number of elements; i.e., there exists no nonempty proper subset 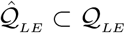 such that 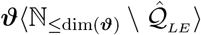 is g.l. identifiable upon fixing the subvector ***ϑ***⟨𝒬 _*LE*_ ⟩;

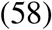

where 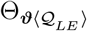 denotes the Cartesian product of the prescribed ranges of the elements of ***ϑ***⟨𝒬_*LE*_ ⟩.

*Proof:* (Statement 1.) Due to Proposition 4.2, we can conclude it by illustrating that the set of all columns of 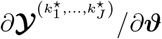 with indices belonging to 𝒬_*l*_, where 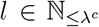, are minimally linearly dependent. This fact can be demonstrated by contradiction. For some 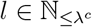, suppose that there exists a nonempty proper subset 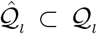 such that the set of all columns of 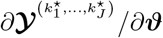 with indices belonging to 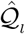 are linearly dependent. Hence, there is a non-identically zero vector 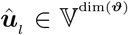, for which 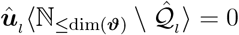, such that

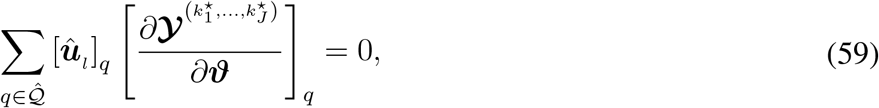

where 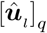 and 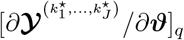 stand for the *q*-th element of ***û*** _*l*_ and the *q*-th column of 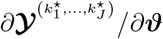, respectively. As a consequence, we have

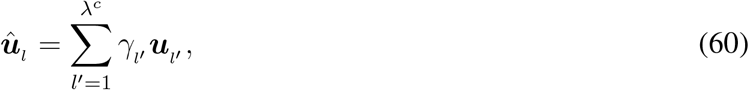

where the scalars *γ*_*l*′_ are meromorphic functions of ***ϑ*** and ***η***_*kwn*_, which are also dependent on ***θ***_*kwn*_ . Since 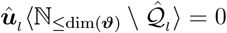 implies ***û***_*l*_ ⟨ℕ_≤dim(***ϑ***)_ \ 𝒬_*l*_ ⟩ = 0, it follows from the RCEF characteristics (57) that all *γ*_*l*′_ except for *l*^′^ = *l* must be zero functions. Thus, we can write ***û***_*l*_ = *γ*_*l*_ ***u***_***l***_ with a non-identically zero function *γ*_*l*_, which contradicts the assumption that 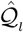 is a nonempty proper subset of 𝒬_*l*_.

(Statement 2.) It can be inferred from the first statement given in (58) and the RCEF characteristics (57) that the set of all columns of 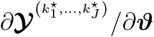 with indices belonging to ℕ_≤dim(***ϑ***)_ \ 𝒬_*LE*_ are linearly independent. Therefore, upon setting the subvector ***ϑ***⟨𝒬_*LE*_ ⟩ to a point in 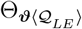 such that 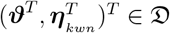, we can ascertain using Test 4.1 that the subvector ***ϑ***⟨ℕ_≤dim(***ϑ***)_ \ 𝒬_*LE*_ ⟩ is g.l. identifiable.

(Statement 3.) According to Test 4.1, it is a direct implication of the fact that the number of the leading entries in ***𝒥***_RCEF_ equals the generic nullity of 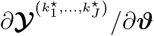.

## V. A priori identifiability of a model for tauopathy in the mouse brain

Here, we first leverage the rich dataset available through the Allen Mouse Brain Connectivity Atlas (AMBCA) [48] (connectivity.brain-map.org) to build an FP-Fisher-KPP Re-Di model for tauopathy progression in the mouse brain. Next, the proposed framework for studying a priori identifiability will be put into practice using this model.

### A. Parameterization on the Allen mouse brain connectome

The inter-region connectivity matrix 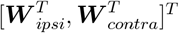 of the Allen Mouse Brain Connectome [48] provides strength estimates of the mutual connections from 213 regions of interest (ROIs) in the right hemisphere to 426 (2 × 213) ROIs in the right (ipsilateral) and left (contralateral) hemispheres. Since the strength values were derived from linear regression, each connection was also assigned a p-value. The matrices ***W*** _*ipsi*_ and ***W*** _*contra*_ are depicted in Fig. 2 for the maximum accepted p-value (significance level) of 0.5.

**Fig. 2:**
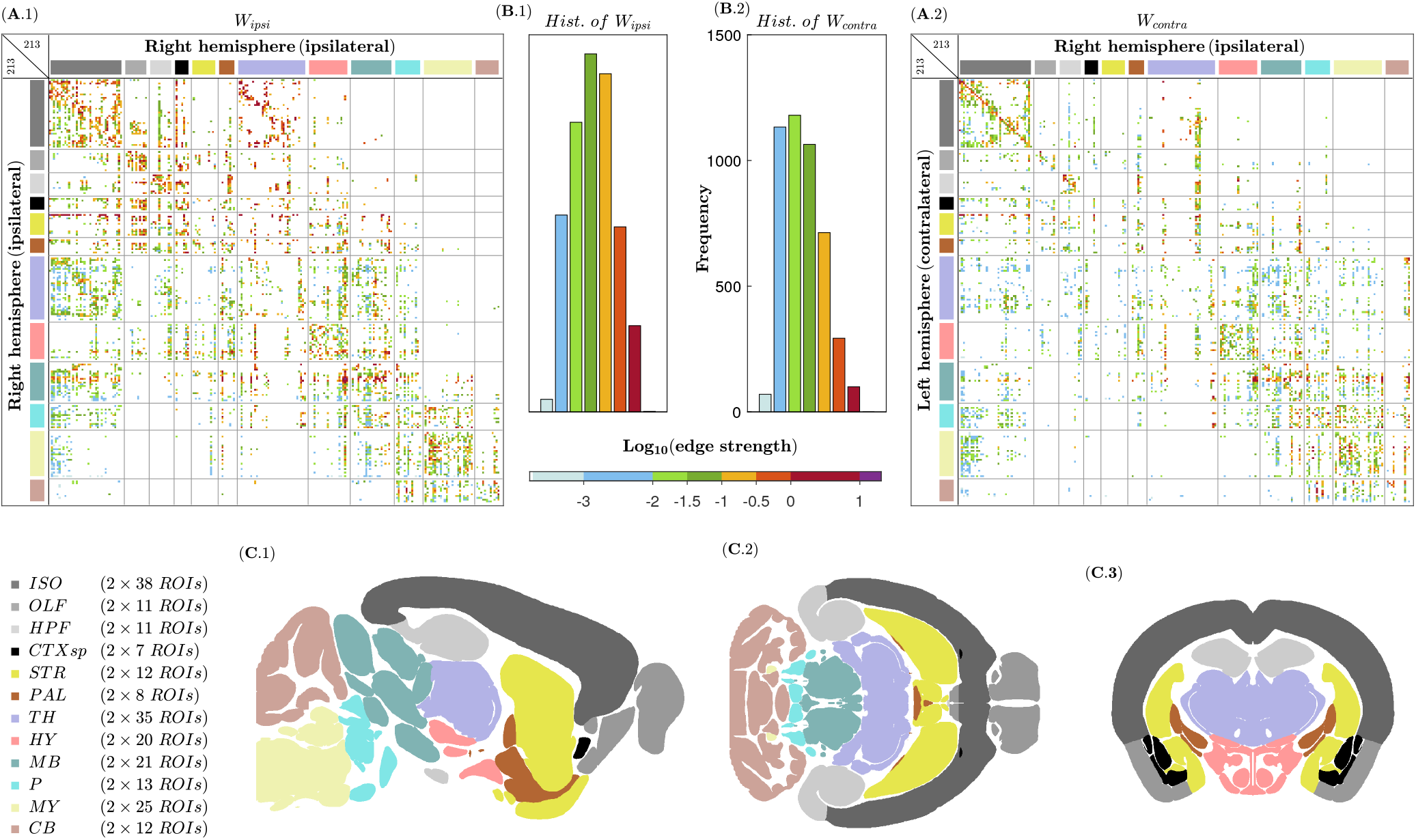
Graphical representations of ***W*** _*ipsi*_ and ***W*** _*contra*_ are shown in (A.1) and (A.2), respectively, at a significance level of 0.5, and their histograms are also displayed in (B.1) and (B.2), respectively. Images (C.1), (C.2), and (C.3) illustrate sagittal, axial, and coronal views of the mouse brain, respectively, depicting the 12 major regions of the gray matter: isocortex (ISO), olfactory areas (OLF), hippocampal formation (HPF), cortical subplate (CTXsp), striatum (STR), pallidum (PAL), thalamus (TH), hypothalamus (HY), midbrain (MB), pons (P), medulla (MY), and cerebellum (CB). To produce this illustration, we used the dataset of the Allen Mouse Common Coordinate Framework (CCFv3) [49] (help.brain map.org/display/mouseconnectivity/API, specifically annotation/ccf 2017).

In the AMBCA project, axonal projections were traced using recombinant adeno-associated virus (AAV) expressing enhanced green fluorescent protein (EGFP), a well-known anterograde tracer. With this understanding, based on the assumption of hemispheric symmetry, one can define the morphological ASA matrix in the following way:

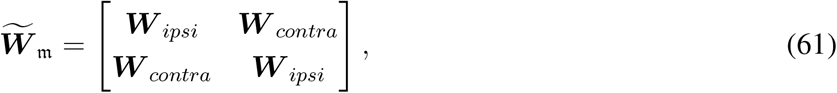

which is not a symmetric matrix. Further, assuming that higher morphological anterograde strength correlates with greater morphological retrograde strength due to both potentially being associated with thicker axonal projections, we can also define the morphological RSA matrix as 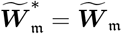.

As shown in Fig. 2, the gray matter can be classified into 12 major divisions, each with different functions, structures, and developmental origins [49]. In this sense, to model the evolution of tau pathology using Eq.(10), one option is to employ a parameterization that is uniformly distributed across each major region, as follows: 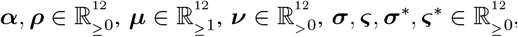, and 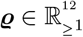 with the same NPD matrix ***B*** ∈ 𝔹^426≥12^, whose (*i, k*)-th element is equal to one if the *i*-th ROI belongs to the *k*-th major region. This parameterization, including 108 (12 × 9) parameters, will be denoted by the pair 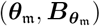, where ***θ***_*m*_ and 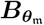 refer to the concatenated parameter vector (12) and concatenated NPD matrix (14), respectively.

Let ***W*** _*m*_ and 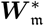 represent the ASA and RSA matrices, respectively, as given by (7), considering the parameterization 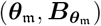, the morphological ASA and RSA matrices 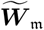 and 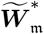. The aim is now to investigate the strong connectivity of the proposed NCB digraph for the mouse brain, or equivalently, the digraph with the weighted adjacency matrix 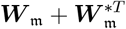 . Let ***A***_gen_ be the generic adjacency matrix derived from the parameterized matrix 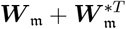 by replacing its non-identically zero elements with one. We say that there is a generic edge from node *j* to node *i* if the (*i, j*)-th element of ***A***_gen_ is equal to one. Consistently, a generic path of length *l* from node *j* to node *i* can also be defined as a sequence of successive generic edges *j* → *j*_2_ → … → *j*_*l*_ → *i*. Using mathematical induction, it can be shown that the number of distinct generic paths of length *l* from node *j* to node *i* is equal to the (*i, j*)-th element of 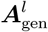. Hence, there exists at least one generic path of length lower than or equal to *l*^⋆^ from each node to every other node if the matrix 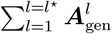 has positive values in all of its off-diagonal elements. Table I reports the minimum values of *l*^⋆^ for different edge-strength thresholds applied to 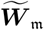.

**TABLE I:**
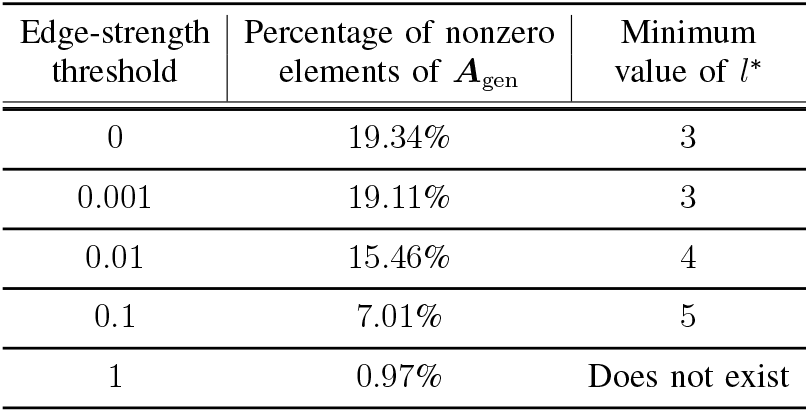
Percentage of nonzero elements of ***A***_gen_ and the minimum values of *l*^*^ for different edge-strength thresholds applied to 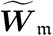.

| Edge-strength threshold | Percentage of nonzero elements of $\mathbf{A}_{\text{gen}}$ | Minimum value of $l^*$ |
| --- | --- | --- |
| 0 | 19.34% | 3 |
| 0.001 | 19.11% | 3 |
| 0.01 | 15.46% | 4 |
| 0.1 | 7.01% | 5 |
| 1 | 0.97% | Does not exist |

As per Table I, the proposed NBC digraph for the mouse brain is strongly connected for almost all values of the diffusion parameter vectors ***σ, ς, σ***^*^, and ***ς***^*^, provided that the edge-strength threshold is at or below 0.1. In the remainder of the manuscript, we will use an edge-strength threshold of 0.01 for all our computation to ensure that the proposed FP-Fisher-KPP Re-Di model for tau pathology propagation possesses Property 3.2 for almost all values of ***σ, ς, σ***^*^, and ***ς***^*^. In fact, the chosen threshold generically guarantees that a non-zero initial concentration at a single arbitrary node can induce non-zero concentrations at all nodes throughout the network over time.

### B. Experimental designs and parameter identifiability

At this point, we will evaluate the parameter identifiability of the proposed FP-Fisher-KPP Re-Di model for tauopathy evolution in the mouse brain across two different experimental scenarios.

In experimental research on tau pathology, PS19 mice, which express the human tau gene with the P301S mutation linked to familial frontotemporal dementia (FTD), serve as valuable animal models for studying the spatiotemporal pattern of tauopathy evolution and its underlying mechanisms [50]. These mice exhibit a nonrandom pattern of tauopathy spreading, beginning in the brainstem and spinal cord around six months of age, and subsequently progressing to the hippocampal formation, including the entorhinal area and hippocampus, as well as the isocortex, covering ROIs such as the somatosensory (SS) and motor (MO) areas [36]. In our first experimental scenario, the parameter identifiability of the proposed model is examined for different designs, each involving a single experiment on PS19 mice. While maintaining the same parameterization 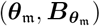 and initial concentration vector 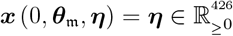 the designs differ in the measured concentration vectors ***y***, resulting in vectors of unknowns ***ϑ*** of varying dimensions, as specified in Table II, and applying Test 4.1 to each of these designs demonstrates that ***ϑ*** is g.l. unidentifiable since the generic rank of 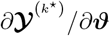 is less than dim (***ϑ***). More preciously, the number of unknowns exceeds the computed generic rank by two for each design; hence, the generic nullity of 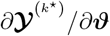 equals 2.

**TABLE II:**
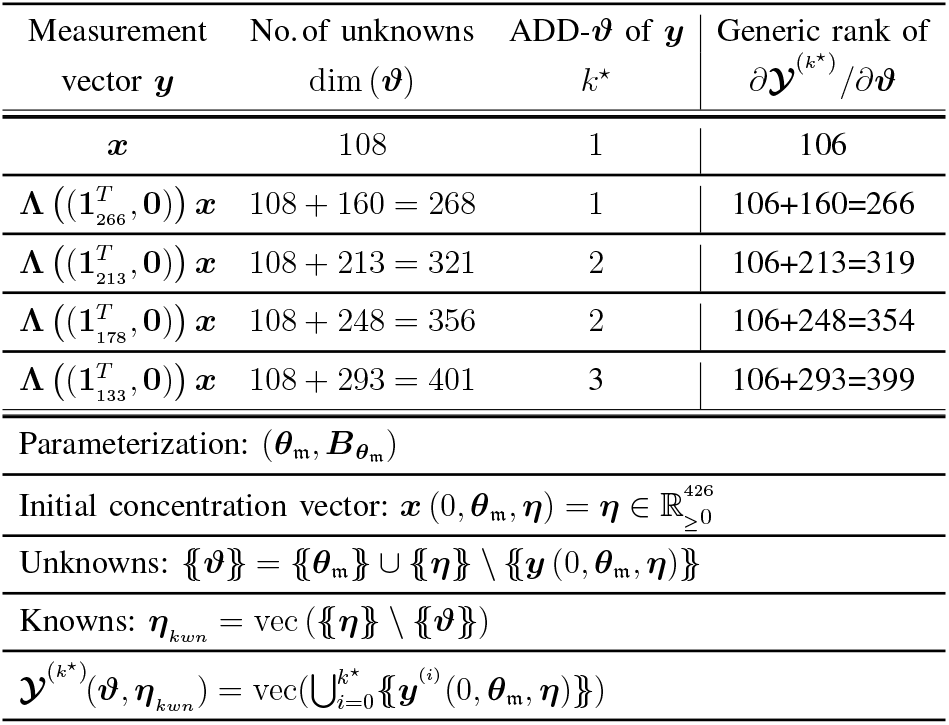
Theoretical designs and symbolic computation results of the first scenario.

Now, let us utilize Proposition 4.3 to determine g.l. minimally dependent unknowns in the design with the measurement vector ***y*** = ***x***. For this design, the rank of 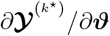 at the randomly selected vectors ***θ***_rs_ and ***η***_rs_, provided in the supplementary PDF file, is equal to its generic rank, i.e., 106. Therefore, since symbolic computations are resource-intensive, we restrict our analysis, without loss of generality, to the point 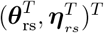 . Numerical calculations yield the matrix ***𝒥***_RCEFm_, as given in (62), which is in RCEF with columns that span the kernel of 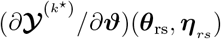.

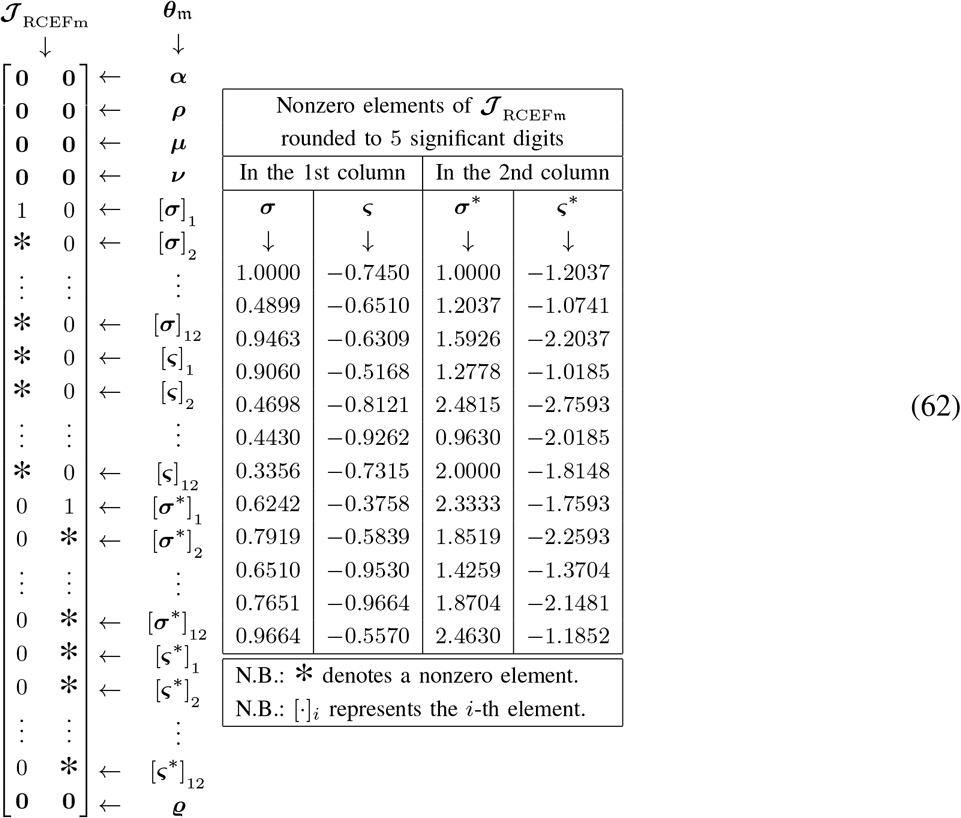

Due to Proposition 4.3, the assessment of ***𝒥*** _RCEFm_ reveals that the elements of the set {{***σ*** }} ∪ {{***ς***}} and those of the set {{***σ***^*^}} ∪ {{***ς***^*^s }} are g.l. minimally dependent, and additionally, upon fixing the elements [***σ***]_1_ and [***σ***^*^]_1_ to almost all of their prescribed values, the remaining unknowns {{***θ***_*m*_ }} \ {[***σ***]_1_, [***σ***^*^]_1_ } become g.l. identifiable. Expanding on this conclusion, it can also be demonstrated that excluding the elements [***σ***]_1_ and [***σ***^*^]_1_ from the vectors of unknowns associated with the other designs make them g.l. identifiable. For each design, the ADD of ***y*** with respect to the newly reduced vector of unknowns remains the same as the one reported in Table II . Since the minimal number of measurement time points for a vector ***y*** to determine a g.l. identifiable vector of unknowns ***ϑ*** is equal to one plus the ADD-***ϑ*** (i.e., the required number of Maclaurin series coefficients) of ***y***, the figures in Table II generally substantiate that a reduction in the number of measurement sites entails an increase in the needed number of measurement time points.

To investigate the in vivo propagation rate of a laboratory-generated tau strain, it is commonly injected into the brain of a transgenic mouse, such as the PS19 [51]. This type of experiment is featured in the second experimental scenario. Specifically, we plan a design that consists of twelve injection experiments whose injection and measurement sites are detailed in Table III . Further, their initial conditions ***x***_*j*_(0, ***θ***_*m*_, ***η***_*j*_) are parameterized by ***H***_*j*_***η***_*j*_ where 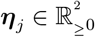, and ***H***_*j*_ ∈ 𝔹^426≥2^ is configured to set the initial concentration at the injection site and at the other ROIs to the first and second elements of ***η***_*j*_, respectively. Assuming that the first elements of the vectors ***σ*** and ***σ***^*^ are predetermined, and given that 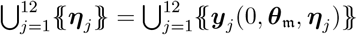, where ***y***_*j*_ (*t*, ***θ***_*m*_, ***η***_*j*_) represents the measured concentration vector from the *j*-th experiment, the vector of unknowns follows as

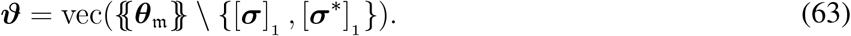

Running Test 4.1 for this design shows that ***ϑ*** is g.l. identifiable, thanks to the fact that the generic rank of *∂****Y*** */∂****ϑ***^(2,…,2)^ is equal to dim (***ϑ***), which is 106, where

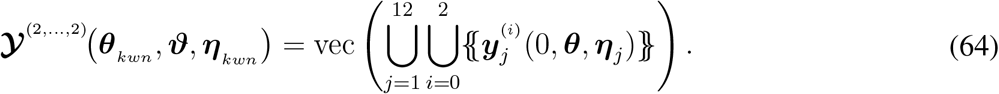

**TABLE III:**
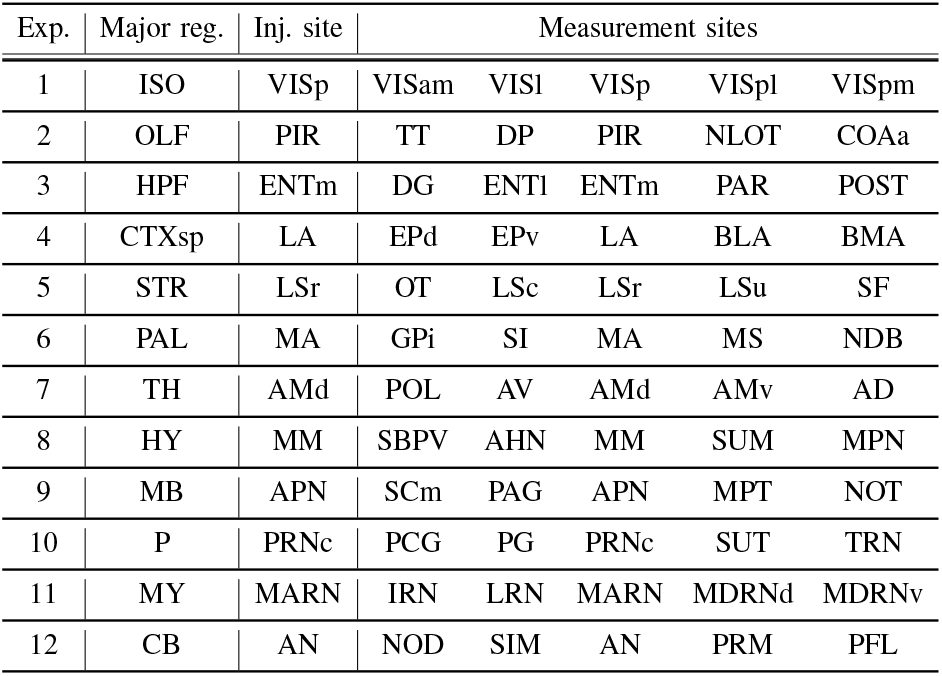
Injection and measurement sites for twelve experiments included in the theoretical design of the second scenario.

## VI. Conclusion

A phenomenological description of tauopathy propagation guided the parameterization of FP-Fisher-KPP Re-Di equations. Afterward, we established our results on generic local identifiability across regular multi-experimental designs for the proposed parameterized model, extending effortlessly to meromorphic systems with regularly parameterized initial conditions. Notably, the concept of generic local minimal dependence was introduced within a Lie group framework, alongside a decomposition method that was developed to further investigate this concept. Ultimately, we constructed a model for tauopathy progression in the mouse brain using the AMBCA dataset, which was then analyzed with the proposed methodology for identifiability analysis.

## Data availability and Supplementary materials

In this research project, we drew on the publicly available data from the Allen Mouse Brain Connectivity Atlas (AMBCA) [48] (connectivity.brain-map.org) and the dataset of the Allen Mouse Common Coordinate Framework (CCFv3) [49] (help.brain-map.org/display/mouseconnectivity/API, specifically annotation/ccf 2017). MATLAB R2022b was also used to carry out all symbolic and numerical computations. Supplementary materials for this article is available in the PDF file named “Supplementary”.

## Supporting information

Randomly selected vectors

## Appendix

### A. Proof of Property 3.2

First, note that the proof for the case of pure diffusion has been presented in Proposition 4.3 of [27]. Before proceeding with the proof, let us rewrite the system (10) as

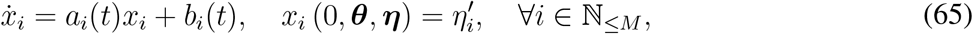

with

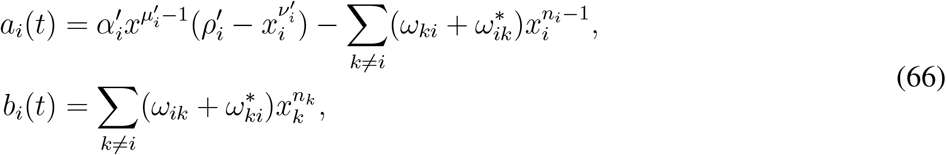

where 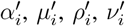, and 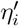 stand for the first element of ***B***_***α***_***α, B***_***µ***_***µ, B***_***ρ***_***ρ, B***_***ν***_***ν***, and ***B***_***η***_***η***, respectively. Referring to the solution formula for a first-order linear differential equation, we get

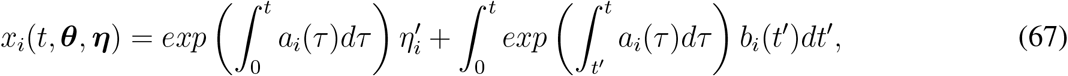

It follows from 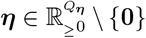 that there is at least one element of ***B η*** that is positive. Let us denote this element by 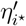. Due to Property 3.1 and the fact that *ω*_*ik*_ ≥ 0 and 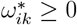 for all *i, k*, the initial condition 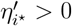 implies 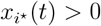 for all *t >* 0 using the formula (67). Since *G* is strongly connected, there is at least one edge from node *i*^⋆^ to another node *l*^⋆^ with a positive anterograde strength or from another node *l*^⋆^ to node *i*^⋆^ with a positive retrograde strength, resulting in 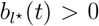 for all *t >* 0. Hence, applying again the formula (67), we deduce that 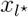 for all *t >* 0. Continuing this reasoning yields 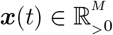 for all *t >* 0.

■

### B. Proof of Lemma (4.1)

Similar to our approach for obtaining 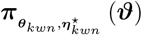 from 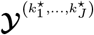, it can be shown that there is a local reparameterization 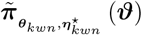 for 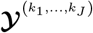. Hence, there locally exist functions 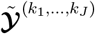 and 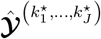 such that

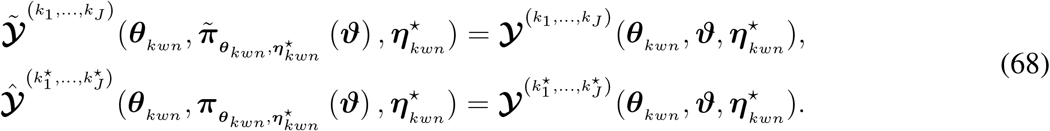

It follows from the equalities in (68) that the ranks of 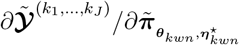 in a neighborhood of 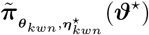 and 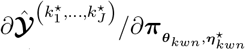 in a neighborhood of 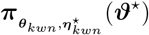 are equal to *λ*. Let 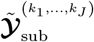 and 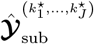denote the subvectors of 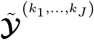 and 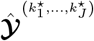, respectively, such that

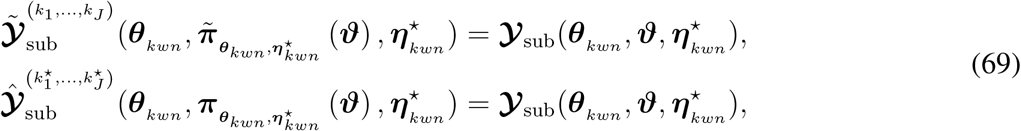

where 𝒴_sub_ comprises all the common elements of 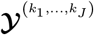 and 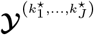. Let us also define the function

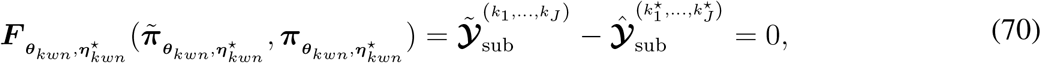

for which the ranks of 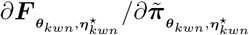 and 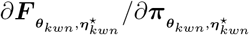 are equal to *λ* in an open subset of its domain containing the point

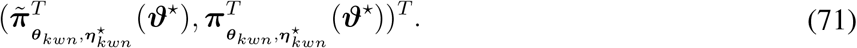

Thus, there is at least one full-rank *λ*-by-*λ* submatrix of 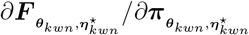 whose corresponding subvector of 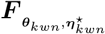 fulfills the conditions of the real analytic implicit function theorem [52], locally ensuring the existence of an analytic function 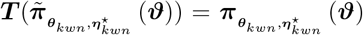. Similarly, the local existence of 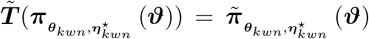 can be concluded. It is evident that, by sufficiently restricting their domains, these functions form a diffeomorphism.

■

