## Supplementary material for "Parameter Dependence in Identifiability Applied to FP-Fisher-KPP Reaction-Diffusion Equations Parameterized for Tauopathy Network Modeling": Randomly selected vectors

Randomly selected vector $\boldsymbol{\theta}_{rs} =$

(1.28, 1.45, 0.57, 1.32, 0.65, 0.89, 0.81, 1.17, 1.19, 1.22, 1.12, 0.97, 1.03, 1.27, 0.95, 1.14, 0.74, 1.35, 1.29, 0.61, 0.58, 1.50, 1.36, 0.89, 2.32, 2.07, 2.09, 2.19, 1.83, 2.49, 2.24, 2.50, 1.85, 1.56, 1.66, 2.09, 1.66, 1.96, 1.85, 1.53, 1.50, 2.29, 2.18, 1.59, 2.02, 2.26, 1.54, 2.36, 1.49, 0.73, 1.41, 1.35, 0.70, 0.66, 0.50, 0.93, 1.18, 0.97, 1.14, 1.44, 1.11, 0.97, 0.94, 0.77, 1.21, 1.38, 1.09, 0.56, 0.87, 1.42, 1.44, 0.83, 0.54, 0.65, 0.86, 0.69, 1.34, 0.52, 1.08, 1.26, 1.00, 0.77, 1.01, 1.33, 0.65, 0.58, 1.19, 0.55, 1.49, 1.09, 0.98, 0.95, 1.22, 0.74, 1.16, 0.64, 2.34, 2.29, 2.22, 2.22, 1.66, 1.92, 2.19, 1.83, 2.21, 1.97, 1.72, 1.73 $)^T$

Randomly selected vector $\boldsymbol{\eta}_{rs} =$

(0.79,0.94,0.97,0.77,0.86,0.78,0.51,0.72,0.82,0.76,0.68,0.96,0.91,0.92,0.68,0.79,0.93,0.96,0.83, 0.60,0.82,0.53,0.70,0.83,0.96,0.90,0.74,0.87,0.70,0.98,0.99,0.93,0.69,0.72,0.62,0.89,0.94,0.95, 0.77,0.79,0.57,0.94,0.72,0.60,0.94,0.88,0.94,0.64,0.83,0.83,0.56,0.70,0.63,0.85,0.64,0.94,0.91, 0.69,0.74,0.84,0.91,0.80,0.78,0.66,0.72,0.85,0.94,0.86,0.50,0.83,0.71,0.71,0.55,0.90,0.66,0.62, 0.67,0.68,0.77,0.78,0.69,0.69,0.75,0.82,0.97,0.86,0.70,0.91,0.56,0.53,0.54,0.58,0.66,0.65,0.50, 0.76,0.54,0.57,0.81,0.92,0.98,0.78,0.99,0.77,0.75,0.66,0.71,0.74,0.53,0.94,0.53,0.71,0.91,0.69, 0.80,0.90,0.94,0.96,0.59,0.62,0.94,0.79,0.75,0.80,0.90,0.76,0.60,0.72,0.71,0.98,0.81,0.84,0.86, 0.67,0.75,0.77,0.57,0.78,0.84,0.71,0.91,0.86,0.68,0.72,0.69,0.88,0.86,0.71,0.84,0.97,0.89,0.85, 0.55,0.69,0.79,0.72,0.52,0.61,0.91,0.50,0.93,0.53,0.83,0.75,0.60,0.78,0.56,0.83,0.79,0.52,0.52, 0.57,0.50,0.71,0.91,0.80,0.76,0.93,0.54,0.95,0.55,0.75,0.57,0.77,0.50,0.88,0.92,0.95,0.99,0.75, 0.63,0.55,0.75,0.79,0.88,0.54,0.83,0.75,0.58,0.96,0.79,0.72,0.97,0.82,0.72,0.91,0.76,0.77,0.84, 0.68,0.61,0.78,0.93,0.70,0.55,0.72,0.65,0.70,0.91,0.70,0.69,0.68,0.57,0.63,0.54,0.71,0.62,0.64, 0.71,0.55,0.74,0.85,0.62,0.89,0.53,0.69,0.50,0.61,0.50,0.59,0.57,0.63,0.58,0.56,0.79,0.95,0.96, 0.61,0.74,0.68,0.76,0.63,0.53,0.71,0.58,0.51,0.97,0.71,0.98,0.88,0.50,0.84,0.85,0.82,0.77,0.60, 0.88,0.61,0.68,0.94,0.92,0.70,0.65,0.80,0.95,0.95,0.79,0.66,0.92,0.72,0.95,0.51,0.76,0.85,0.58, 0.66,0.59,0.66,0.70,0.77,0.52,0.77,0.63,0.62,0.62,0.57,0.97,0.96,0.90,0.86,0.58,0.68,0.59,0.50, 0.65,0.84,0.81,0.77,0.71,0.64,0.75,0.88,0.88,0.78,0.87,0.82,0.56,0.75,0.67,0.54,0.57,0.59,0.83, 0.71,0.84,0.62,0.50,0.76,0.63,0.97,0.95,0.69,0.51,0.83,0.91,0.98,0.52,0.72,0.79,0.84,0.85,0.82, 0.86,0.68,0.79,0.55,0.52,0.98,0.64,0.79,0.98,0.59,0.59,0.67,0.96,0.69,0.63,0.57,0.69,0.68,0.56, 0.71,0.54,0.80,0.50,0.78,0.89,0.61,0.72,0.78,0.53,0.74,0.82,0.61,0.91,0.98,0.92,0.75,0.63,0.87, 0.61,0.97,0.81,0.80,0.58,0.54,0.62,0.92,0.95,0.84,0.86,0.61,0.78,0.90,0.70,0.99,0.54,0.66,0.75, 0.53,0.86,0.77,0.76,0.91,0.92,0.89,0.65,0.72,0.87,0.55,0.55,0.63,0.76,0.98,0.85,0.65,0.64,0.92, 0.95,0.81,0.62,0.54,0.91,0.79,0.97,0.53$)^T$
